# A segment of *Triticum timopheevii* chromosome 3G confers type II Fusarium head blight resistance and reduces DON accumulation in wheat

**DOI:** 10.1101/2025.10.27.684833

**Authors:** Andrew Steed, Surbhi Grewal, Roshani Badgami, Julie King, Ian P King, Paul Nicholson

**Author notes:** These authors contributed equally to this work and share first authorship (Andrew Steed, Surbhi Grewal).

## Abstract

Fusarium head blight (FHB) is a major disease of small grain cereals that is particularly damaging on wheat and can result in yield losses of up to 80%. Resistance to FHB in wheat is broadly classified as resistance to initial infection (type I) or resistance to disease spread within the spike (type II). A high level of type I FHB resistance was reported in an accession of wheat wild relative *Triticum timopheevii*. Hexaploid wheat-*T. timopheevii* introgression lines carrying a segment of the short arm of chromosome 3G (Chr3G) of this accession expressed high levels of FHB resistance following spray inoculation. Further analyses of these introgression lines showed that the Chr3G segment confers a potent type II resistance, accompanied by improved grain weight retention, and reduced deoxynivalenol (DON) accumulation in grain. These results indicate that the Chr3G resistance has the potential to dramatically reduce FHB susceptibility and DON accumulation in grain in wheat cultivars. An introgression of a segment of Chr7A^t^ into the short arm of chromosome Chr7A also enhanced type II FHB resistance.

## INTRODUCTION

FHB is a major disease of both hexaploid and tetraploid wheat, with plants most susceptible to infection during flowering. Furthermore, *Fusarium graminearum sensu stricto*, *Fusarium culmorum*, and *Fusarium asiaticum* produce trichothecene mycotoxins such as deoxynivalenol (DON), which is toxic to both humans and animals. Improving resistance to FHB has proved to be a very challenging target for plant breeders. While more than 250 QTL for FHB resistance have been identified in hexaploid wheat (Jia et al., 2018), only a small number have been defined as single genes and shown to be potent and robust. One of the best characterised of these is *Fhb1*, located on chromosome 3B (Chr3B), which has been attributed to a histidine-rich calcium-binding protein (Li et al., 2019; Su et al., 2019).

Introgressions from wild relatives have had a major impact on wheat production and disease resistance. An example of this is the *Aegilops ventricosa* 2NS introgression, which confers a yield advantage, as well as being the major source of wheat blast resistance. The 2NS introgression is present in over 85% of CIMMYT wheat varieties and represents the sole effective source of wheat blast resistance (Kishii, 2019). Several of the major FHB resistance genes/QTL reported have origins in wild relatives (Mawcha et al., 2022). FHB resistance is generally classified into two broad types: resistance to initial infection (type I) and resistance to disease spread in the spike (type II) (Schroeder and Christensen, 1963). The major effect QTL *Fhb3* originates from *Leymus racemosus* and has been transferred to Chr7A of wheat, where it confers mainly type II resistance (Zhu et al., 2019). *Fhb6* originates from *Elymus tsukushiensis* and has been transferred into the short arm of Chr1A of wheat, again conferring a high level of type II resistance (Cainong et al., 2016). *Fhb7* is one of only two cloned FHB resistance genes, the other being *Fhb1* (Wang et al., 2020; Li et al., 2019; Su et al., 2019). Several alleles of *Fhb7* have been reported originating from *Thinopyrum elongatum* (Wu et al., 2024) and *Thinopyrum ponticum* (Zhang et al., 2022), and they confer potent type II resistance when introgressed into wheat (Wu et al., 2024). Type II FHB resistance has also recently been reported in wheat lines generated by introgression of the long arm of Chr3S^c^ from *Reogneria ciliaris* and addition lines carrying Chr4Ns from *Psathyrostachys huashanica* (Song et al., 2024; Li et al., 2025). These reports highlight the potential of wheat relatives as donors of FHB resistance to wheat.

A previous report (Steed *et al.,* 2022) identified two *T. timopheevii-*bread wheat introgressions, with differing sizes of the same region of Chr3G, conferring a potent FHB resistance upon spray inoculation. These authors also noted that the donor *T. timopheevii* accession possessed a very high level of type I resistance but appeared to lack type II resistance. Given the lack of type II resistance in the *T. timopheevii* donor, it was assumed that the resistance exhibited in introgression lines following spray inoculation was a result of type I resistance. In this paper we describe the characterisation of introgression lines for type II resistance and demonstrate that introgression of the segment of Chr3G confers potent type II resistance in hexaploid wheat, improved grain weight retention, and reduced DON accumulation in grain. An introgression of a segment of Chr7A^t^ into chromosome Chr7A also enhanced type II FHB resistance but to a lesser extent.

## MATERIALS AND METHODS

### Plant materials

The wheat–*T. timopheevii* introgression lines analysed in this study were described previously by King et al. (2022). Wheat cv. Paragon and additional introgression lines (BC2F4-40, BC4F2-112) were obtained from the Wheat Research Centre (University of Nottingham). The latter were generated following the same crossing and backcrossing strategy described in King et al. (2022). All plants were grown under standard glasshouse conditions with a 16 h photoperiod at 25 °C.

### Whole-genome sequencing and analysis

Genomic DNA for skim-sequencing was extracted from 10-day-old leaf material using the protocol of Grewal et al. (2022), with the addition of a phenol–chloroform purification step to improve DNA quality. Library preparation and sequencing were carried out by Novogene (UK) Company Limited. Each library was skim-sequenced to ∼0.05× coverage on a NovaSeq 6000 S4 flow cell with paired-end 150 bp reads (Coombes et al., 2023).

Sequence data were analysed using the pipeline described by King et al. (2025). To assess the introgressed *T. timopheevii* genomic regions, reads were aligned to a combined reference assembly comprising wheat cv. Chinese Spring RefSeq v1.0 (IWGSC et al., 2018) and the *T. timopheevii* genome (Grewal et al., 2024). Per-base read depths were aggregated into 1 Mb non-overlapping bins. For wheat chromosomes, coverage in each sample was normalised by dividing bin counts by the corresponding median-smoothed values from the Paragon control. For *T. timopheevii* chromosomes, coverage was normalised within each sample by scaling bin counts to the median of bins above a minimum coverage threshold, providing a relative measure of dosage independent of the wheat control. Normalised values were capped at 2× for visualisation. Contiguous stretches of bins with reduced or elevated coverage were interpreted as candidate introgressions.

### FHB disease assessment of *T. timopheevii* introgression lines by point inoculation (2022, 2023)

Lines carrying *T. timopheevii* introgressions and the hexaploid susceptible recipient wheat variety (Paragon) were assessed for FHB resistance following point inoculation in two sets of experiments (2022, 2023). Between 8 and 51 individual spikes per line from multiple plants were point-inoculated at mid-anthesis in the 2022 trial and between 10 and 50 in the 2023 trial. Inoculum (10 µl) of a DON-producing *F. graminearum* isolate (1×10^6^ conidia ml^-1^) was introduced directly into a central spikelet for each spike. Disease was assessed at intervals post-inoculation according to the rate of symptom development and the number of infected spikelets above and below the point of inoculation recorded.

### Grain weight analysis

Spikes from five plants of Paragon and each introgression line were harvested at maturity. The grain from each spike was separated by position relative to the point of inoculation: above and below. The ‘above’ and ‘below’ components from the infected spikes from each plant were combined to produce each sample, with plants acting as replicates. For each line, four representative non-inoculated ears were sampled and split above and below the region where treated plants were inoculated. Grain number and grain weight were determined for above and below the point of inoculation for treated and control plants. The hundred grain weight of treated spikes was calculated as a percentage of the control for each sampled plant. These values are referred to as relative grain weight (RGW).

### DON analysis

DON content of grain above and below the point of inoculation was quantified. Grain for 3 of the 5 replicate samples used for estimation of RGW was ground to a fine powder in a pestle and mortar. DON was quantified using AgraStrip® Pro WATEX® lateral flow devices according to the manufacturer’s recommendations.

The Extraction buffer bag provided in the AgraStrip® Pro WATEX® test kit was dissolved in 50 ml of distilled water and shaken vigorously just before use. All procedures were adjusted to account for the weight of the sample relative to that used in routine analysis. Flour (1 g) of each sample was weighed out and added to a 15 ml Falcon centrifuge tube. Extraction buffer (5 ml) was added to each tube, and the contents were shaken for 2 minutes. Following this, tubes were centrifuged at 2000 x g for 1 minute. The AgraVision™ Pro reader machine was set up using the quick guide procedures. Dilution Buffer (1 ml) was added to an Eppendorf tube, and supernatant (100 µl) of sample extract was added and mixed. The Eppendorf was centrifuged (2000 x g) for 30 seconds. The AgraStrip Pro Deoxynivalenol WATEX lateral flow cartridge was inserted into the port of the AgraVision™ Pro reader. Diluted extract (200 µl) was added to the lateral flow cartridge, and results were read from the machine following analysis.

### Statistical analysis

All statistical analyses were performed using Genstat 18th. Disease assessment trials were blocked, and general linear model (GLM) was used to calculate predicted means and standard errors for infected spikelets above and below the point of infection with “Block”, “Inoculation date” and “Line” included in the model. Assessment of RGW and DON concentration used GLM to calculate predicted means and standard errors with “Replicate” and “Line” included in the model.

## RESULTS

### Characterisation of *T. timopheevii* Chr3G introgressions in wheat

Skim-sequencing was used to characterise a panel of 18 wheat–*T. timopheevii* introgression lines alongside the recurrent wheat parent Paragon, providing precise boundaries of introgressed segments at the level of 1 Mb bins (**Supplemental Table S1**). This analysis refined and corrected the earlier chromosome-specific KASP marker-based descriptions (Grewal et al., 2020; King et al., 2022). Four small introgressions (<40 Mb) not previously detected were identified in intervals between adjacent KASP markers. Conversely, some introgressions reported earlier were absent, likely reflecting spurious KASP signals. In contrast, others were reinterpreted as deletions in the wheat background that had shifted marker clustering in the earlier study.

Although the panel contained introgressions from several *T. timopheevii* chromosomes, eleven lines carried segments of Chr3G. Previously, Steed et al. (2022) had implicated Chr3G in conferring FHB resistance. Therefore, these lines, together with wheat control Paragon, are highlighted in **Figure 1**. Paragon displayed generally uniform coverage across wheat Chr3B and Chr3D, with no evidence of large-scale *T. timopheevii* Chr3G segments. However, skim-sequencing revealed a previously unidentified ∼7 Mb fragment from the distal end of Chr3GS on wheat Chr3DS (**Supplemental Table S1**), indicated by an increase in read coverage on the short arm from Chr3G and a drop in read coverage on the short arm of Chr3D (**Figure 1**). As this fragment appears fixed in the wheat background and is potentially from a different *T. timopheevii* accession introduced from an ancestral introgression event, it was treated as a background feature rather than a candidate resistance interval.

**Figure 1.**
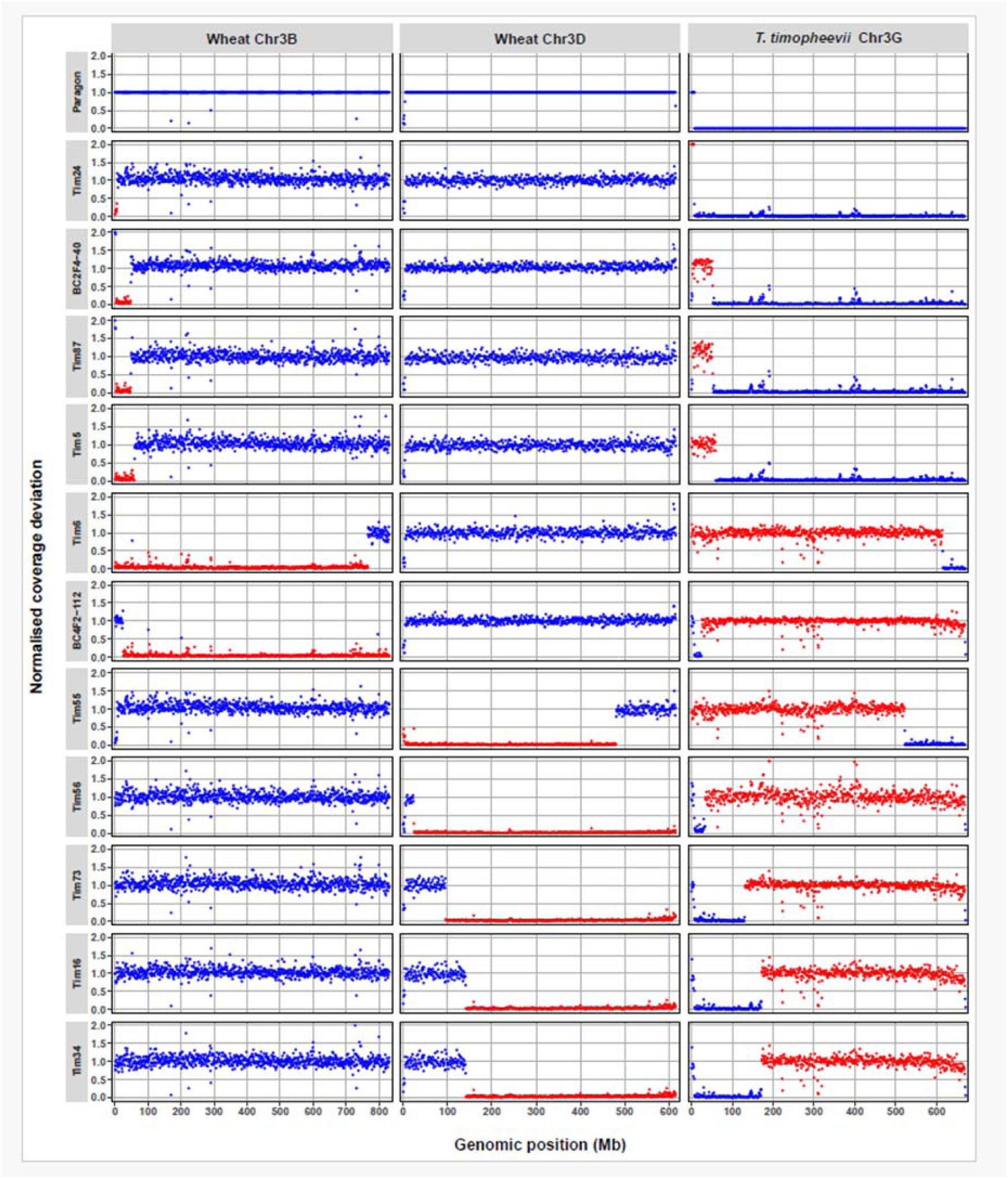
Coverage deviation profiles across homoeologous group 3 chromosomes in wheat cv. Paragon and wheat-*T. timopheevi*i introgression lines. Normalised read coverage (y-axis) was calculated in 1 Mb bins (x-axis) along wheat Chr3B and Chr3D, and *T. timopheevii* Chr3G. Blue points indicate background coverage, while red points highlight contiguous regions of coverage deviation that represent putative introgressions. Each row corresponds to one line, with Paragon included as the control used for normalisation of reads. Values are capped at 2× for display.

Among the introgression lines, reciprocal patterns of reduced coverage on wheat Chr3B or Chr3D together with elevated coverage on Chr3G confirmed clear chromosomal substitutions. Tim87 and BC2F4-40, like Tim5, carried a small Chr3G introgression at the distal end of Chr3BS, whereas Tim6 and BC4F2-112 harboured extensive Chr3G substitutions spanning much of Chr3B. Tim16, Tim34, Tim55, Tim56 and Tim73 carried large Chr3G introgressions replacing wheat Chr3D. Several lines (BC4F2-112, Tim16, Tim56 and Tim73) also retained the ∼7 Mb background Chr3GS fragment detected in Paragon (**Figure 1).**

By expanding beyond the previously characterised resistant lines Tim5 and Tim6 (Steed et al., 2022), which first implicated Chr3G in FHB resistance, this study assembled a broader panel of recombinants carrying different portions of Chr3G. This provides the foundation for comparing resistant and susceptible lines and progressively narrowing the candidate interval of *T. timopheevii* Chr3G responsible for resistance.

### Assessment of type II FHB resistance in wheat*-T. timopheevii* introgression lines (2022)

In 2022, fifteen wheat*-T. timopheevii* introgression lines, together with the recurrent wheat parent Paragon, were assessed for type II FHB resistance in a polytunnel environment. Eight of these lines carried *T. timopheevii* Chr3G segments, either replacing regions of wheat Chr3B or Chr3D (**Figure 1**). The panel in this study also includes lines with *T. timopheevii* introgressions from other chromosomes, and lines with Chr3G segments not previously associated with resistance in the study of Steed et al. (2022) (**Supplemental Table S1**). All lines were evaluated by point inoculation to measure disease spread within the spike above and below the point of inoculation. Significant differences in disease spread were observed among lines with some exhibiting extremely restricted disease development. Examples of FHB symptoms on spikes of Paragon and Tim6 four weeks after inoculation are shown in **Figure 2**.

**Figure 2.**
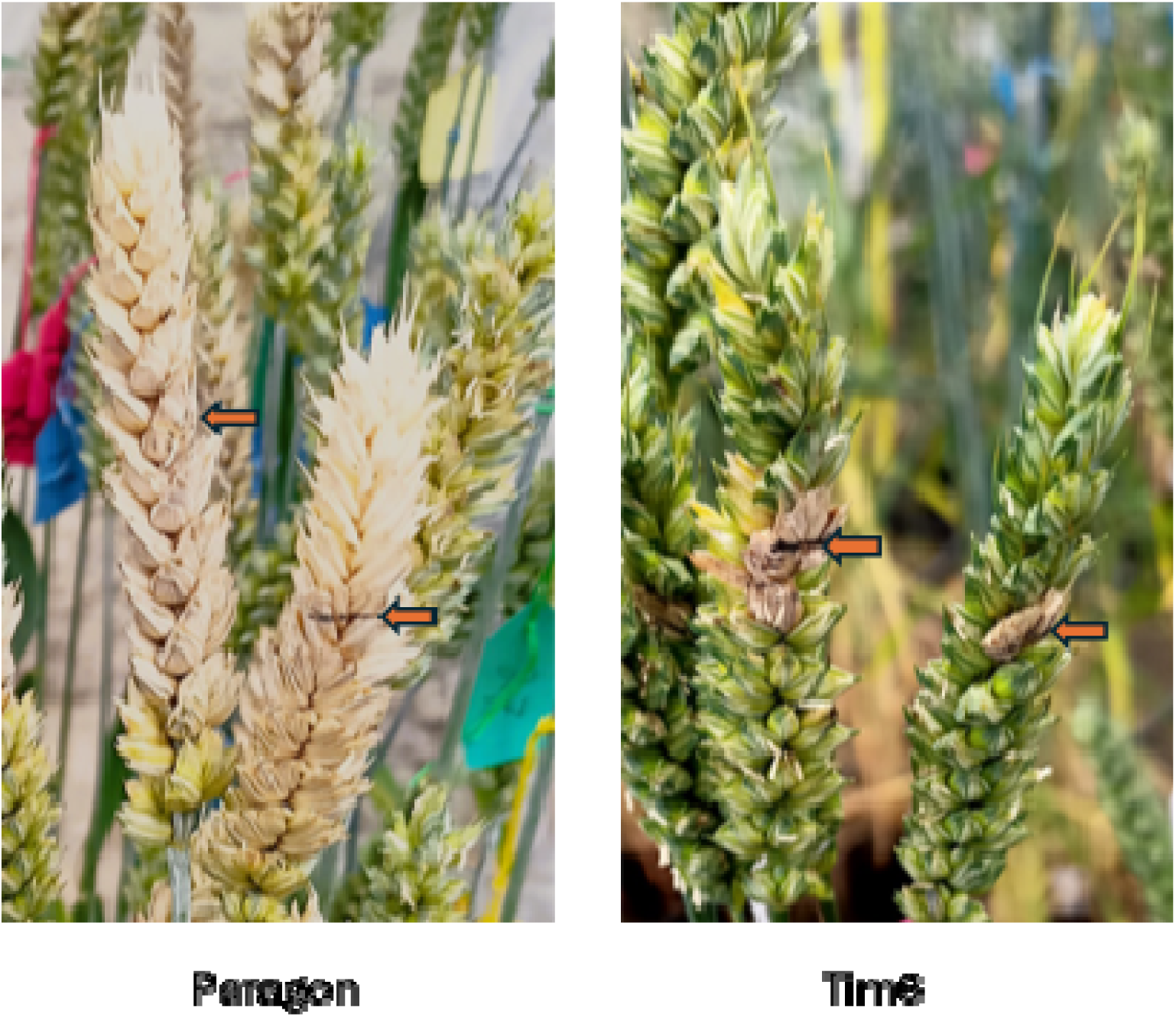
Representative examples of FHB symptoms on spikes of Paragon parent line and introgression line Tim6 carrying a Chr3G segment following inoculation of the central spikelets with conidia of *F. graminearum*. The inoculated spikelets are indicated by arrows.

Disease spread was assessed both above and below the point of inoculation, and results from 12 days post-inoculation (dpi) are shown (**Figures 3a, 3b**). Six lines exhibited significantly less disease spread (Tim5, Tim6, Tim11, Tim33, Tim55 and BC2F4-40) both above and below the point of inoculation than the recipient line Paragon. Three of these lines (Tim5, Tim6, and BC2F4-40) contained introgressions, including the Chr3G region identified in previous work (Steed et al., 2022). The fourth highly resistant line (Tim55) carried a large Chr3G substitution replacing most of wheat Chr3D (**Figure 1**). These results indicate that several Chr3G-containing introgressions (whether on Chr3B or Chr3D) are associated with enhanced resistance to FHB spread in the spike.

**Figure 3.**
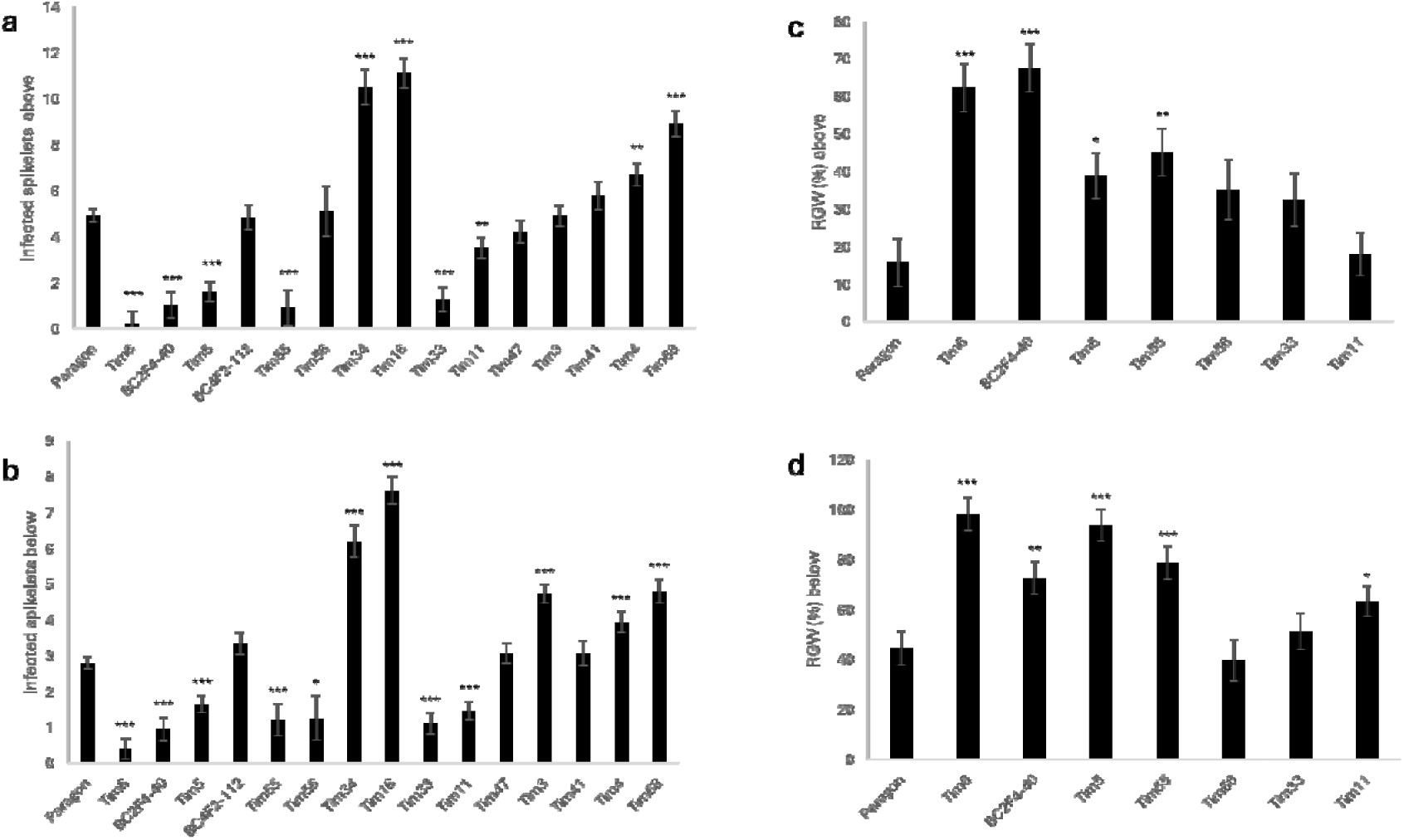
(a) Disease spread above point of inoculation at 12 days post-inoculation (dpi). (b) Disease spread below the point of inoculation (12 dpi). (c) Predicted mean for calculated hundred grain weight of infected spikelets above the point of inoculation as a percentage compared to uninfected spikes. (d) Predicted mean for calculated hundred grain weight of infected spikelets compared to uninfected spikelets below the point of inoculation as a percentage compared to uninfected spikes. Relative grain weight, RGW.

Previous work (Steed et al., 2022) also identified other *T. timopheevii* introgressions that enhanced resistance following spray inoculation. Lines Tim11 and Tim33 had similar levels of resistance as lines containing Chr3G introgressions. Tim11 contains a short segment of Chr6A^t^ (501-549 Mb) while Tim33 contains a large segment of Chr7A^t^ (0-225 Mb). Following point inoculation, Tim11 and Tim33 exhibited slower symptom development than Paragon, but both sustained similar loss in grain weight as Paragon by the time of harvest (**Figures 3c, 3d**). Two additional lines carrying segments of Chr7A^t^ were also assessed in an attempt to refine the location of the Chr7A^t^ FHB resistance. Tim47 carries two segments of Chr7A^t^ (47-98 and 112-548 Mb) while Tim68 has a segment from 45-130 Mb (**Supplemental Table S1**). Neither line showed enhanced FHB type II resistance, with Tim68 being significantly more susceptible than Paragon (**Figures 3a, 3b**).

Two lines (BC4F2-112 and Tim56) showed levels of disease spread similar to Paragon above and below the point of inoculation and were less resistant than lines carrying smaller Chr3G introgressions on Chr3B (**Figures 3a, 3b**). While BC4F2-112 carried a large Chr3G introgression on Chr3B, Tim56 carried a large Chr3G introgression on Chr3D (**Figure 1**). Line Tim3 showed significantly greater disease spread than Paragon only below the point of inoculation, while Tim41 had levels of disease similar to Paragon. Four lines (Tim4, Tim16, Tim34, and Tim68) had significantly greater disease spread than Paragon both above and below the point of inoculation (**Figures 3a, 3b**). Two of these lines (Tim16 and Tim34) contained large *T. timopheevii* Chr3G substitutions on wheat chromosome Chr3D that extended towards the telomere of the long arm. However, these Chr3G introgressions differed in size and position from those in resistant lines such as Tim5 and Tim6, suggesting that the regions associated with resistance and susceptibility are distinct.

### Grain weight retention following infection (2022)

Spikes were harvested from a selection of lines, and grain was separated into above and below the point of inoculation. Grain weight of inoculated spikes was determined relative to grain from non-inoculated spikes (relative grain weight, RGW) (**Figures 3c, 3d**). The four type II FHB resistant lines (Tim5, Tim6, Tim55 and BC2F4-40) showed significantly greater RGW than Paragon. Tim56, which carries a large Chr3G introgression on Chr3D, did not differ from Paragon for RGW above or below the point of inoculation (**Figures 3c, 3d**). Tim11 and Tim33 did not show sustained improvements in RGW compared to Paragon.

### Assessment of type II FHB resistance in wheat*-T. timopheevii* introgression lines (2023)

Type II FHB resistance was subsequently assessed in a different subset of introgression lines during early spring 2023 using a controlled-environment cabinet. The panel included several lines tested previously and three additional lines (Tim35, Tim73 and Tim87). Disease spread above the point of inoculation at 21 dpi was significantly lower in all six lines carrying Chr3G introgressions (Tim5, Tim6, Tim55, Tim56, Tim87, and BC2F4-40) than in Paragon (**Figure 4a**). Disease spread below the point of inoculation was also significantly lower than in Paragon for five of these six lines (**Figure 4b**), with the exception of Tim56, which did not differ significantly from the control and carries a large Chr3G introgression on Chr3D extending along the long arm (**Figure 1**). BC4F2-112, which contains a large Chr3G segment on Chr3B, showed a level of disease spread similar to Paragon both above and below the point of inoculation (**Figures 4a, 4b**). Tim35, which carries small introgressions on Chr1A and Chr7B, also showed susceptibility comparable to Paragon, while Tim73, containing a Chr3G introgression that replaces the long arm of Chr3D, was significantly more susceptible than Paragon (**Figures 4a, 4b**).

**Figure 4.**
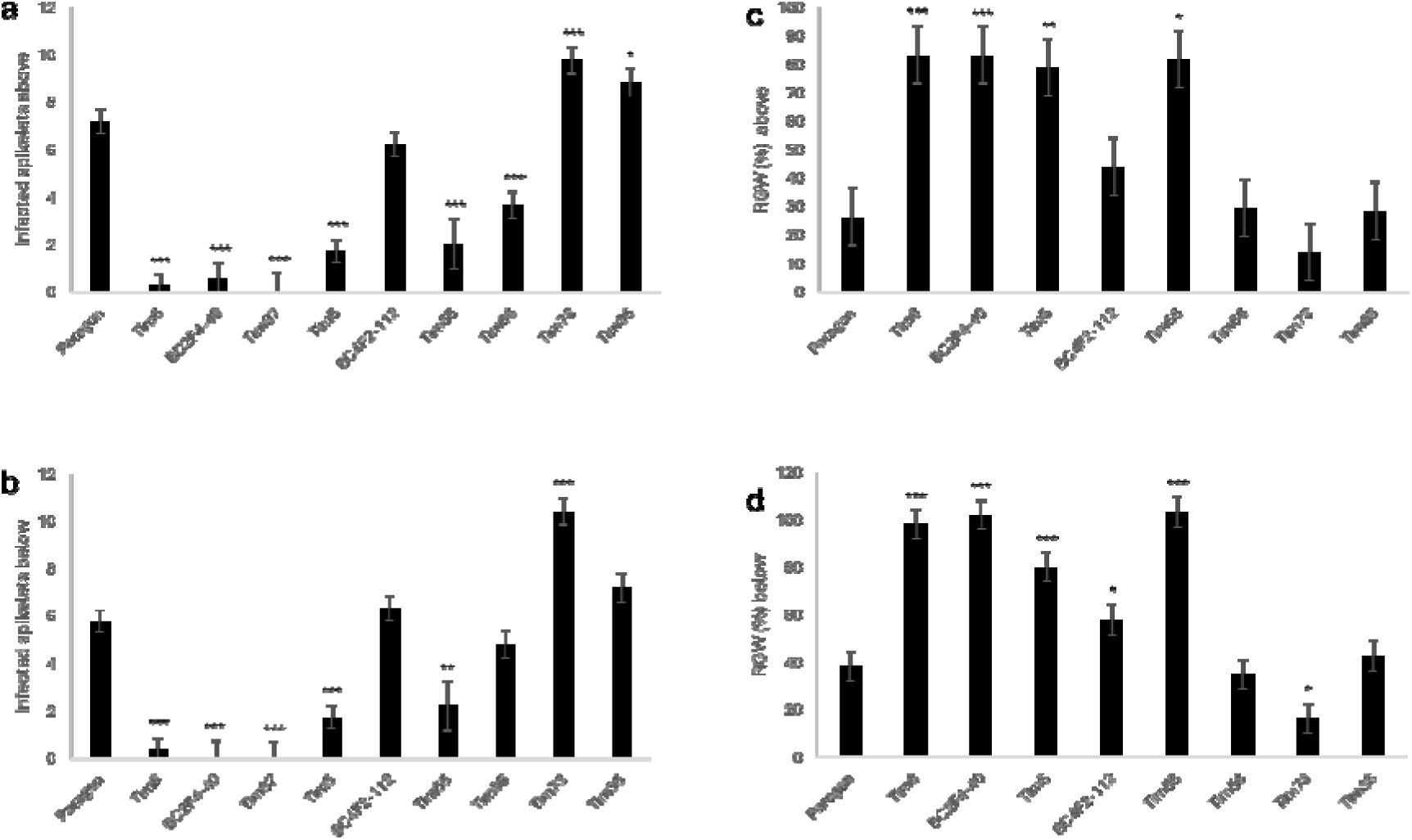
(a) Disease assessment of infected spikelets above the point of inoculation 21 days post-inoculation (dpi). (b) Disease assessment of infected spikelets below the point of inoculation 21 dpi. (c) Predicted mean for calculated hundred grain weight as a percentage of the control above the point of inoculation. (d) Predicted mean for calculated hundred grain weight as a percentage of the control below the point of inoculation.

### Grain weight retention following infection (2023)

Sufficient spikes and grain were available to determine RGW for all lines except Tim87. The four lines carrying Chr3G introgressions previously associated with increased resistance (Tim5, Tim6, Tim55 and BC2F4-40) all had significantly greater RGW than Paragon both above and below the point of inoculation (**Figures 4c, 4d**). Three of the other lines (Tim35, Tim56 and BC4F2-112) exhibited similar losses in RGW to Paragon both above and below the point of inoculation (**Figures 4c, 4d**). RGW below the point of inoculation of line Tim73 was significantly lower than that of Paragon. This line also exhibited greater disease spread than Paragon, which is likely reflected in the reduced RGW (**Figures 4a, 4b**).

### DON accumulation in infected grain (2023)

One of the most important factors in FHB disease is the accumulation of trichothecene mycotoxins such as deoxynivalenol (DON) in the grain of infected plants (Amarasinghe et al., 2019). There was sufficient grain from inoculated spikes below the point of inoculation to quantify DON content (**Figure 5**). The level of DON in Paragon was 266 mg/kg. Comparable levels were detected in Tim35 (191 mg/kg), which contains only the background *T. timopheevii* segment found in Paragon, and in Tim56 (179 mg/kg), which carries a large Chr3G introgression on Chr3D. The four lines that displayed the highest levels of type II resistance in disease and grain weight assessments (Tim5, Tim6, Tim55 and BC2F4-40) all accumulated substantially less DON than Paragon (23, 0.2, 3.5 and 1.2 mg/kg, respectively). In contrast, the highly susceptible line Tim73 showed markedly elevated DON content (626 mg/kg). These findings are consistent with the observed disease severity and RGW across lines, indicating that lines with reduced FHB spread also exhibited reduced toxin accumulation.

**Figure 5.**
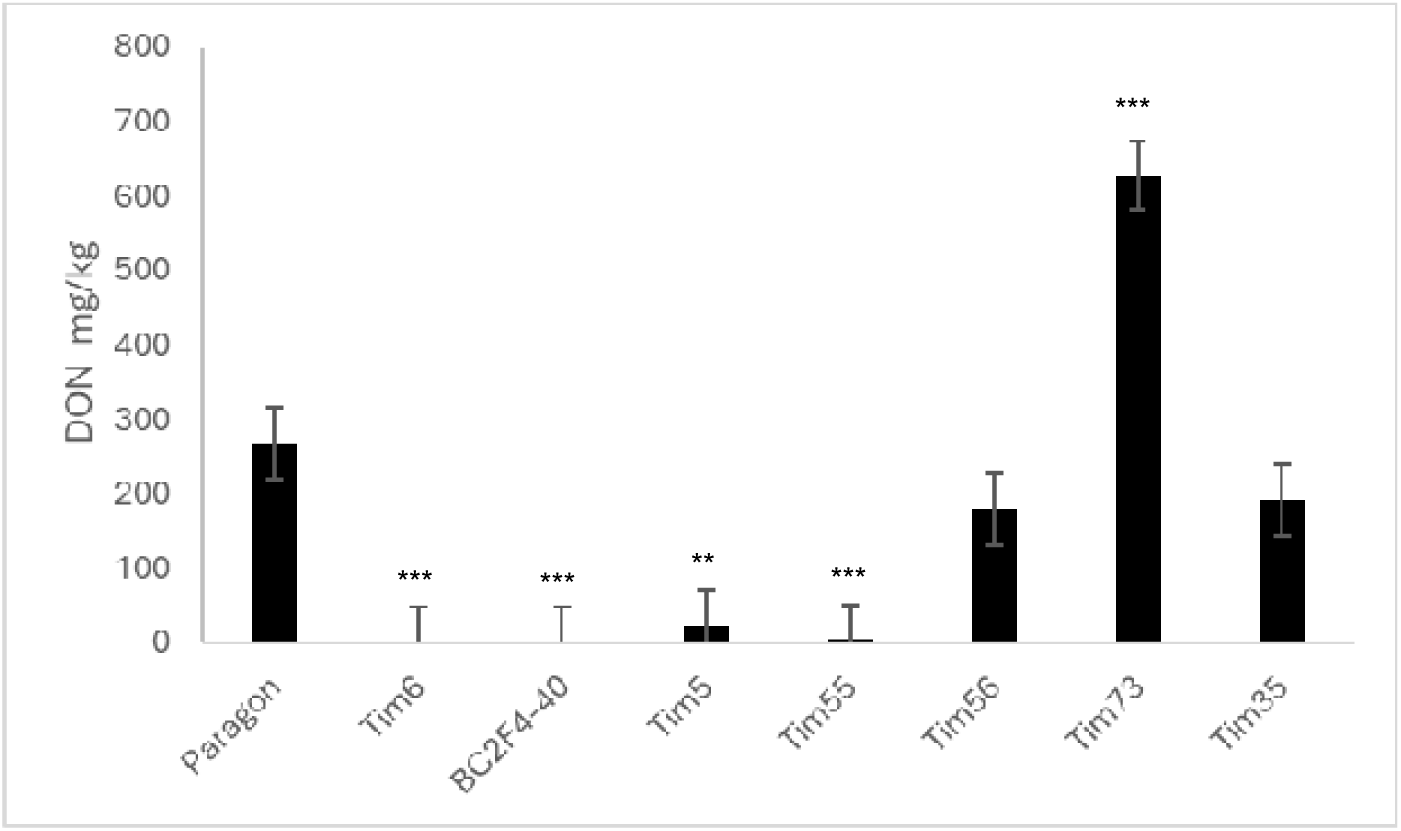
Deoxynivalenol (DON) content (mg/kg) of harvested grain from spikelets below the point of inoculation in Paragon and Paragon-timopheevii introgression lines.

### Mapping the genomic interval associated with type II FHB resistance on Chr3G

The FHB responses of the wheat-*T. timopheevii* introgression lines to point inoculation with *F. graminearum* were compared with the size and position of their Chr3G introgressions and the wheat chromosome into which they were introduced. This comparison revealed that the potent type II resistance derived from *T. timopheevii* is associated with a distal short-arm interval on Chr3G extending from approximately 3 to 25 Mb (**Figure 6**). Lines carrying this interval, irrespective of whether it replaced part of Chr3B or Chr3D, consistently exhibited high levels of resistance, reduced disease spread, and low DON accumulation. The lower boundary of this interval is defined by line BC2F4-40, which is resistant despite lacking 0–3 Mb of Chr3G, whereas the upper boundary is supported by BC4F2-112 and Tim56, both of which are non-resistant and contain Chr3G in the region beyond 25 Mb.

**Figure 6.**
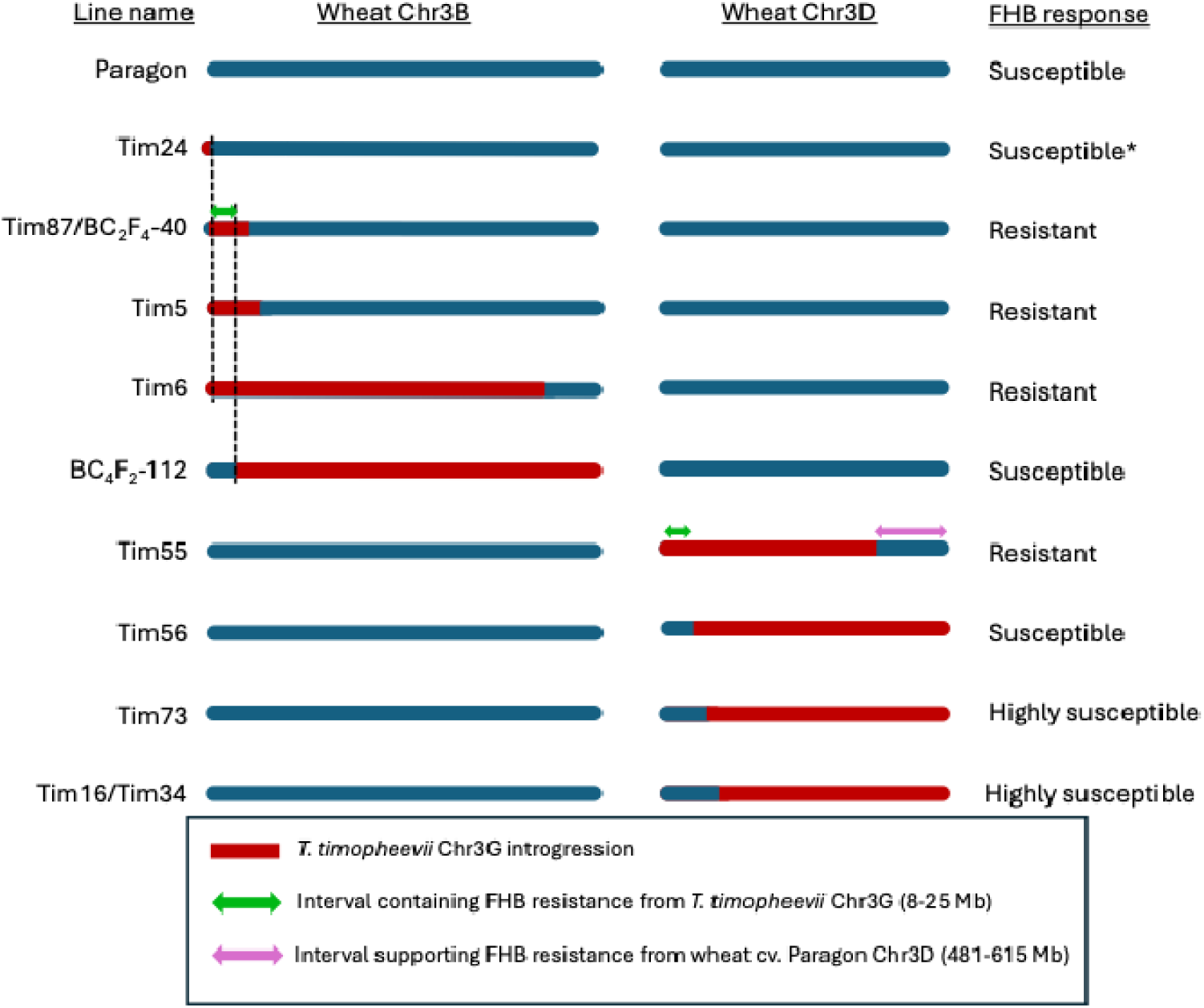
Representation of the wheat Chr3B and Chr3D in Paragon and Paragon-*Triticum timopheevii* introgression lines. Wheat DNA is represented in blue and *T. timopheevii* DNA is represented in red. The regions associated with FHB resistance originating from *T. timopheevii* Chr3G and Paragon Chr3D are indicated by the green and purple double-headed arrows on Chr3B and Chr3D respectively. * = Susceptibility determined in Steed et al (2022).

Lines in which Chr3G introgressions did not include the 3–25 Mb interval showed little or no improvement in resistance relative to Paragon. The effect of Chr3G introgressions on Chr3D was also influenced by the extent of replacement of the distal long arm of Chr3D. Lines such as Tim16, Tim34 and Tim73, which lacked this distal region, displayed greatly enhanced susceptibility. This contrasts with lines carrying large Chr3G substitutions on Chr3B, such as BC4F2-112, which did not show increased susceptibility despite lacking the 3–25 Mb interval. These observations suggest that the distal portion of Chr3DL in Paragon contributes to FHB resistance, and that loss of this region increases susceptibility unless accompanied by the *T. timopheevii* segment spanning 3–25 Mb on Chr3G (**Figure 6**). One exception, Tim56, which carries a large Chr3G introgression on Chr3D extending across the long arm but lacking the 3–25 Mb resistance interval, was moderately FHB susceptible, similar to Paragon rather than exhibiting the high level of susceptibility seen in other Chr3D-substitution lines. This may indicate compensation from other introgressed loci from *T. timopheevii*, such as the Chr7A^t^ segment.

While the precise boundaries cannot be resolved beyond the 1 Mb binning used for this analysis, the results indicate that *T. timopheevii*-derived type II FHB resistance lies within a discrete interval on Chr3GS between roughly 3 and 25 Mb, and that loss of the distal region of Paragon Chr3D increases susceptibility unless compensated by this *T. timopheevii* segment.

## DISCUSSION

### Previously reported sources of FHB resistance in *Triticum timopheevii*

Resistance to FHB has been reported in several studies involving *T. timopheevii*. In an early report, one of three accessions was found to exhibit moderate type II resistance while all three lacked resistance to initial infection (type I) (Yong-Fang et al., 1997). Accession PI 343447 of *T. timopheevii* was used to generate the FHB resistant spring wheat line TC 67 that was then crossed to the susceptible variety Brio. Characterisation of 230 F_7_ recombinant inbred lines for type II FHB resistance under glasshouse and field conditions identified two QTL for FHB resistance (Malihipour et al., 2017). Both QTL were identified on the long arm of Chr5A, with one near the centromere in the interval between markers cfd6.1 and barc48 and the second more distal between cfd39 and cfa2185. The more potent and distal QTL was associated with reduced grain shrivelling (Fusarium Diseased Kernels) in field trials and reduced severity in glasshouse trials following point inoculation, indicating that it conferred type II resistance (Malihipour et al., 2017).

Two QTL, contributing type II resistance to FHB, were identified in a separate population developed between wheat line PI 277012 that includes *T. timopheevii* in its pedigree and wheat variety Grandin (Chu et al., 2011). Both QTL were also located on chromosome Chr5A, with one on the short arm and one on the long arm. The QTL on the short arm is located in a similar position to *Qfhs.ifa-5A* identified in Sumai 3, with both being in the region of marker XBarc180 (Chu et al., 2011). The QTL on the long arm corresponds to the same region as the potent locus identified in TC 67, derived from *T. timopheevii* PI 343447, making it highly likely that these represent the same resistance (Chu et al., 2011). The FHB resistance QTL on Chr5A probably derives from Chr5A^t^ of *T. timopheevii* as the A^t^ genome is more closely related to the A genome of wheat than the G genome of *T. timopheevii* (King et al., 2022; Grewal et al., 2024).

### Mapping and genetic basis of the 3G-associated resistance

It was previously reported that accession PI 94760 (referred to as Tim_P95-99.1-1 by Steed et al., 2022) possessed a very high level of type I FHB resistance but lacked appreciable levels of type II resistance. In that study, several wheat–*T. timopheevii* introgression lines carrying segments of the distal portion of Chr3GS on Chr3B exhibited greatly increased resistance following spray inoculation, consistent with type I resistance. In the present work, fine comparison of introgression structure and type II FHB response delimited the resistance to an interval on the short arm of Chr3G between approximately 3 and 25 Mb (**Figure 6**). Sequencing confirmed that Tim24 carries a small (∼8 Mb) segment from the distal end of Chr3GS on Chr3B in addition to the background Chr3G fragment present on Chr3D (**Figure 1**). Although this line was not phenotyped for type II resistance in the present study, its previously reported susceptibility following spray inoculation (Steed et al., 2022) indicates that the lower boundary of the resistance interval may lie near 8 Mb. The absence of type II resistance in the donor accession despite the presence of this interval suggested that its expression might depend on interactions with the wheat genomic background, a hypothesis explored further below.

Whole-genome sequencing revealed that Paragon contains a previously unidentified ∼5 Mb region at the distal end of Chr3DS that is highly similar to an equivalent region of ∼7 Mb on Chr3GS (**Supplemental Table S1**). This was evident from increased read coverage on Chr3G accompanied by a corresponding reduction on Chr3D (**Figure 1**). As this segment is fixed in the Paragon background, it likely represents a historical recombination event between wheat and a tetraploid wild ancestor, possibly *T. timopheevii* or a closely related species. With the advent of extensive genome and exome sequencing, such relic introgressions are increasingly being recognised as evidence of ancient gene flow from wild relatives that contributed to the adaptive diversity of modern bread wheat (He et al., 2019; Heuberger et al., 2024).

Based on KASP marker data, lines Tim2, Tim24 and Tim35 were originally reported to contain small segments (up to 10.75 Mb) of Chr3G introgressed onto the distal end of Chr3B (King et al., 2022). None of these lines exhibited enhanced FHB resistance following spray inoculation, and it was inferred that the FHB resistance locus on Chr3G lay proximal to this region (Steed et al., 2022). In the present study, Tim35 was also found to lack type II resistance. Whole-genome sequencing revealed that Tim35 carries a ∼100 Mb deletion on

Chr3B, which caused the KASP assay to falsely indicate the presence of Chr3G. The small Chr3G-like signal detected actually corresponds to the background Chr3G segment present on Chr3D in Paragon (**Supplemental Table S1**). Sequencing also confirmed that Tim24 carries a small (∼8 Mb) segment from the distal end of Chr3GS on Chr3B, in addition to the background Chr3G fragment present on Chr3D (**Figure 1**). Although Tim24 was not assessed for type II resistance in the present work, it was found to be susceptible following spray inoculation (Steed et al., 2022). Spray inoculation assesses the combined effects of type I and type II resistances, while point inoculation only assesses levels of type II resistance. Since Tim24 was susceptible to spray inoculation, it is reasonable, though not conclusive, to assume that Tim24 would also be susceptible to point inoculation. If so, the resistance-associated interval would lie between approximately 8 and 25 Mb on Chr3GS (**Figure 6**).

The introgressed region on Chr3G associated with enhanced type II FHB resistance contains approximately 328 high-confidence genes according to the *T. timopheevii* genome annotation (Grewal et al., 2024). A diverse range of genes implicated in disease resistance have been characterised in the Triticeae (Li et al., 2025), many of which encode proteins containing leucine-rich repeat (LRR) and/or kinase domains. Of the 47 disease resistance genes cloned in wheat, 32 encode nucleotide-binding leucine-rich repeat (NLR) proteins (Wulff and Krattinger, 2022). Four NLR genes (Tritim_EIv0.3_0555910, _0559470, _0560200, and _0560430) are located within the resistance-associated interval and could potentially contribute to the observed FHB resistance phenotype. In addition to NLRs, the 8–25 Mb region contains 18 other genes predicted to encode LRR-containing proteins, four of which also possess kinase domains (Tritim_EIv0.3_0554450, _0554550, _0557780, and _0561540). Such LRR receptor-like kinases (LRR-RLKs) play central roles in the recognition of pathogen-associated molecular patterns (PAMPs), triggering PAMP-triggered immunity (PTI) (Ding et al., 2024). LRR-RLKs are abundant across plant genomes, with 223 identified in *Arabidopsis* and 309 in rice, and their functions extend beyond defence to regulation of growth and development. For example, the product of *BRASSINOSTEROID-INSENSITIVE 1* (*BRI1*) perceives brassinosteroids that regulate cell elongation and developmental processes (Guo et al., 2013). Interestingly, mutation of barley *BRI1* enhanced resistance to *Fusarium culmorum* while reducing growth rate (Goddard et al., 2014), highlighting a potential trade-off between growth and defence that may also be relevant here.

Beyond these LRR-related genes, 54 additional genes within the interval encode proteins predicted to contain kinase domains of diverse types. Given the number and diversity of potential defence-related gene candidates in this interval, extensive functional analysis will be required to identify which specific gene or genes underlie the type II FHB resistance derived from *T. timopheevii*.

### Interactions between 3G and 3D loci influencing type II FHB resistance

Susceptibility factors to FHB have been reported previously where loss of a portion of the wheat genome or replacement with a chromosome from another species led to reduced susceptibility to FHB (Hales et al., 2020; Chhabra et al., 2021). Thus, the resistance observed following introgression of a segment of Chr3G replacing its equivalent region of Chr3B might reflect the loss of an FHB susceptibility factor rather than the gain of a resistance determinant. However, the enhanced resistance associated with introgression of this section of Chr3G was also observed when the introgression replaced the corresponding region of Chr3D rather than Chr3B, suggesting that resistance is conferred by the addition of resistance factor(s) on Chr3G rather than the removal of susceptibility factors from wheat.

Several of the lines tested (Tim16, Tim34 and Tim73) carried large Chr3G substitutions replacing much of the long arm of Chr3D and showed markedly greater susceptibility to FHB than Paragon, with extensive disease spread and elevated DON accumulation (**Figures 3 and 4**). In contrast, Tim55, which carries a large Chr3G substitution on Chr3D but retains the distal region of Chr3DL, exhibited a high level of resistance. These observations suggest that loci within the distal Chr3DL interval (approximately 481–615 Mb) may contribute to type II FHB resistance in Paragon and that their absence in lines where Chr3DL is entirely replaced by Chr3GL increases susceptibility. Notably, all highly resistant lines retained Chr3DL, supporting the interpretation that Chr3GS-mediated resistance functions optimally in the presence of Chr3DL. Although no line combines a Chr3GS introgression with loss of Chr3DL, the available evidence implies that expression of Chr3GS-derived resistance depends on this distal 3DL region.

The *T. timopheevii* donor accession (PI 94760) itself does not exhibit type II resistance, despite carrying the complete Chr3G chromosome. This suggests that the Chr3GS resistance determinant is not inherently inactive but rather requires interaction with factors present in the wheat genome. The amphiploid (wheat × *T. timopheevii*) line described by Steed et al. (2022), which carries both chromosomes Chr3G and Chr3D, displayed strong type II resistance, supporting the hypothesis that expression of Chr3GS resistance depends on a permissive interaction with loci on Chr3DL. In contrast, when Chr3DL is entirely replaced by Chr3GL, as in Tim16, Tim34 and Tim73, this interaction is lost, leading to enhanced susceptibility.

The relationship between Chr3GS, Chr3GL and Chr3DL is further illustrated by the behaviour of BC4F2-112 and Tim56. Both lines lack the Chr3GS resistance interval yet contain the entire long arm of Chr3G and are only moderately susceptible, not severely so. This pattern indicates that the presence of Chr3GL alone does not drive susceptibility and that the extreme susceptibility observed in lines such as Tim16, Tim34 and Tim73 is most likely due to the complete loss of the distal Chr3DL region rather than an inhibitory effect of Chr3GL. In Tim56, the large Chr7A^t^ introgression may further mitigate susceptibility, an aspect explored later, where Chr7A^t^-derived contributions to FHB resistance are examined.

Taken together, these results indicate that Chr3GS provides the principal resistance determinant (between 8-25 Mb), whose expression depends on a supportive factor located on the distal region of Chr3DL (between 481-615 Mb). Loss of this Chr3DL region, or its replacement by Chr3GL, disrupts the interaction and increases susceptibility, whereas retention of Chr3DL allows full expression of resistance. This model accounts for the high resistance of Tim5, Tim6, Tim55 and the amphiploid, the severe susceptibility of lines lacking Chr3DL, and the moderate susceptibility of lines such as Tim56 and BC4F2-112. It also provides a mechanistic explanation for the absence of type II resistance in *T. timopheevii* itself, which carries Chr3GS but lacks the complementary Chr3DL factor present in wheat. QTL on Chr3D for type I, type II FHB resistance and DON accumulation have been reported in several studies (Wu et al 2022; Cai et al 2019). The meta-analysis of Chinese wheat landraces identified a QTL on the long arm of Chr3D associated with type II FHB resistance and it is conceivable that the effect of the replacement of Chr3DL with Chr3GL reflects loss of the more positive allele (Cai et al 2019).

### Additional FHB resistance associated with chromosome 7A□

In addition to lines carrying segments of Chr3G, Tim11 (Chr6A^t^: 501-549 Mb) and Tim33 (Chr7A^t^: 0-225 Mb), were reported to have greatly enhanced resistance following spray inoculation (type I resistance) (Steed et al 2022). Lines Tim11 and Tim33 had similar levels of type I resistance as lines containing Chr3GS. In the current study of type II resistance, both lines exhibited slower symptom development than Paragon but sustained a similar loss in grain weight as Paragon by the time of harvest (**Figure 3**). These results indicate that the resistance conferred by the Chr6A^t^ and Chr7A^t^ segments from *T. timopheevii* functions more as a type I (resistance to initial infection) rather than type II resistance (resistance to spread in the spike). A similar differential effect has been reported for *Fhb5* (synonym for *Qfhs.ifa-5A*) that contributes more towards type I resistance than to type II resistance (Buerstmayr et al., 2003).

In addition to the FHB resistance conferred by the 8-25 Mb region of Chr3G, a region of chromosome Chr7ALJ between approximately 42 and 128 Mb was previously reported to be associated with increased resistance (Steed et al., 2022). Five additional lines carrying Chr7ALJ introgressions were examined in the present study to refine this interval. Skim sequencing revealed that Tim3 and Tim4 each carry a single Chr7ALJ segment spanning 119-545 Mb and 0-9 Mb, respectively. Tim47 carries two Chr7ALJ segments (47-98 Mb and 112-548 Mb), while Tim68 has a single segment from 45–130 Mb (**Supplemental Table S1**). Tim41 was not subjected to skim sequencing, but earlier KASP marker analysis indicated that it contains a segment of Chr7ALJ extending from the distal end of the short arm to at least 3.6 Mb but not beyond 41 Mb (King et al., 2022). None of these lines exhibited enhanced type II FHB resistance, and Tim68 was significantly more susceptible than Paragon (**Figure 3**).

Comparison of the Chr7ALJ introgressions and FHB responses across Tim33, Tim41, Tim47, and Tim68 suggests that the enhanced resistance previously observed in Tim33 (Steed et al., 2022) may be conferred by gene(s) located within the 9-45 Mb region of Chr7ALJ. Introgression of a segment from the short arm of chromosome Chr7LLJ of *Leymus racemosus* into wheat chromosome Chr7A has also been reported to enhance resistance, and the corresponding locus was designated *Fhb3* (Qi et al., 2008). It may therefore not be coincidental that introgressions from two different wild relatives (*T. timopheevii* and *L. racemosus*) into the short arm of Chr7A both improve resistance, even if acting primarily at different stages of infection. The short arm of Chr7A has been reported to carry susceptibility factor(s) to FHB (Chhabra et al., 2021), so replacement of this wheat segment with those from *T. timopheevii* or *L. racemosus* may lead to increased resistance through removal of susceptibility factors rather than the introduction of novel resistance genes.

Line Tim56 lacks both the resistance-associated interval on the short arm of Chr3G (8-25 Mb) and the region of Paragon Chr3DL (481-615 Mb) associated with FHB resistance (**Figure 1**). However, Tim56 was not significantly more susceptible than Paragon in any of the parameters evaluated across assays (**Figures 3 and 4**). It is assumed that this line does not exhibit the very high level of FHB susceptibility seen in Tim16, Tim34 and Tim73 because of the contribution of the Chr7ALJ segment it carries (0-225 Mb), which may partially compensate through resistance gene(s) located within the 9-45 Mb interval implicated in Tim33.

### Implications for breeding and future applications

In this study, *T. timopheevii* Chr3G was shown to confer a high level of type II FHB resistance when introgressed into hexaploid wheat cv. Paragon. The resistance was accompanied by enhanced grain weight retention and markedly reduced DON accumulation in infected grain. Given that DON contamination is one of the main determinants of economic loss and food safety concern in cereals, these results are of considerable applied importance. The financial impact of downgrading wheat from human to animal feed due to mycotoxin contamination has been estimated at approximately €3 billion across Europe over the past decade (Johns et al., 2022). Incorporation of the Chr3G segment into elite cultivars could therefore provide a means to improve both disease resilience and grain quality.

This study also reinforces the critical role that genetic diversity from wild relatives plays in improving wheat resilience. With advances in genomics and molecular cytogenetics, it is now feasible to systematically transfer beneficial alleles from ancient progenitors into modern cultivars and to track these introgressions using diagnostic DNA markers to support marker-assisted selection (Othmeni et al. 2025).

While numerous QTL for FHB resistance have been identified in hexaploid wheat (Jia et al., 2018), only a limited number have been detected in durum wheat (Zhao et al., 2018). Durum wheat remains generally more susceptible to FHB, and resistance transfer from bread wheat has proven challenging (Buerstmayr et al., 2012). It has been proposed that critical resistance genes may reside on the D genome, absent in tetraploid wheat, or that durum wheat harbours suppressor loci that inhibit resistance expression (Ban and Watanabe, 2001; Kishii et al., 2005). Future work should therefore aim to introgress and evaluate *T. timopheevii* Chr3G segments in durum wheat backgrounds to determine whether they can similarly confer FHB resistance.

## DATA AVAILABILITY

Raw skim-sequence reads for wheat-*T. timopheevii* introgression lines have been deposited at the European Nucleotide Archive (ENA) under project accession PRJEB6515.

## AUTHOR CONTRIBUTIONS

AS, PN, SG, JK, IPK: Conceptualization and manuscript writing. PN, JK, IK, SG: funding acquisition. SG, IK and JK: germplasm generation. SG: sequence data analysis. AS, RB and PN carried out the disease screening of plants. AS: disease data analysis. All authors have read and agreed to the published version.

## ACKNOWLEDGEMENTS

AS and PN thank the Horticultural Services team of the John Innes Centre for their input into care of the plants used in these experiments. In particular, we would like to thank Lewis Hollingsworth for his assistance throughout the duration of this project.

